# Historical biogeography and phylogeography of *Indoplanorbis exustus*

**DOI:** 10.1101/2021.05.28.446081

**Authors:** Maitreya Sil, Juveriya Mahveen, Abhishikta Roy, K. Praveen Karanth, Neelavara Ananthram Aravind

## Abstract

The history of a lineage is intertwined with the history of the landscape it resides in. Here we showcase how the geo-tectonic and climatic evolution in South Asia and surrounding landmasses have shaped the biogeographic history of *Indoplanorbis exustus*, a tropical Asian, freshwater, pulmonated snail. We amplified partial COI gene fragment from all over India and combined this with a larger dataset from South and Southeast Asia to carry out phylogenetic reconstruction, species delimitation analysis, and population genetic analyses. Two nuclear genes were also amplified from one individual per putative species to carry out divergence dating and ancestral area reconstruction analyses. The results suggest that *Indoplanorbis* dispersed out of Africa into India during Eocene. Furthermore, molecular data suggests *Indoplanorbis* is a species complex consisting of multiple putative species. The primary diversification took place in Northern Indian plains or the Northeast India. The speciation events appear to be primarily allopatric caused by a series of aridification events starting from late Miocene to early Pleistocene. None of the species seemed to have any underlying genetic structure suggestive of high vagility. All the species underwent population fluctuations during the Pleistocene likely driven by the Quaternary climatic fluctuations.

## Introduction

Diversity of a biogeographic subregion is maintained through dispersal and diversification. Much of the vertebrate fauna of the Indian subcontinent was assembled through dispersal (Karanth 2021). However, their origins are extremely diverse, owing to the strategic position of the subcontinent (Mani, 1974). Furthermore, lineages adapted to a range of habitats and terrains could successfully colonize the subcontinent because of its geographic and climatic heterogeneity. The climate also fluctuated over time, which allowed for dispersal of both wet adapted and dry adapted groups into and out of India (Klaus et al., 2016, Sil et al., 2019). The aforementioned climatic fluctuations over time as well as the relative isolation of the subcontinent also facilitated in-situ diversifications (Karanth, 2015; Agarwal and Karanth, 2015; Agarwal and Ramakrishnan, 2017; Deepak and Karanth, 2017; Lajmi and Karanth, 2020). However, the studies are too sparse to develop any generalities and largely focuses on terrestrial vertebrates.

The ecology, physiology, dispersal ability of freshwater organisms is vastly different and as a result they respond differently to similar climatic changes or geographic features (Kappes et al., 2014; Lundberg et al., 1998). Hence, it is imperative to understand how diversity of freshwater organisms is maintained in a region in order to obtain a holistic idea about the evolutionary history of fauna of a region. Freshwater snails are unique in terms of their mode of dispersal among freshwater organisms in that they disperse along a river basin with the flow of current, but also across basins, when they attach themselves to feathers or feet of aquatic birds (Kappes and Haase, 2012; Van Leeuwen et al., 2013). Hence, these taxa could give us unique insight into how the physical features of the subcontinent affect distribution, diversification, and phylogeography in the freshwater ecosystems. *Indoplanorbis exustus* is a pulmonate freshwater snail belonging to family Bulinidae. *Indoplanorbis* is a monotypic genus distributed across tropical Asia from Sundaland, throughout mainland Southeast Asia, parts of China, all over Indian subcontinent, till Iran in the West. They were also introduced in West Asia, Africa and French West Indies. The habitat for these air-breathing snails include all kinds of water-bodies stagnant parts of lotic and lentic, including seasonal ephemeral bodies of water. However, they are not found in high elevation and fast flowing streams. They serve as intermediate hosts to *Schistosoma indicum* group of parasitic trematodes which cause schistosomasis in cattle and can also cause cercarial dermatitis in humans (Agarwal et al., 2000; Morgan et al., 2002; Wooi, Chiew Eng Lim and Ambu, 2007). The family Bulinidae is distributed in Africa and Asia. The sister genus *Bulinus* is found throughout Africa and Madagascar. Several hypotheses were put forward to explain the colonization of India by the lineage leading to *Indoplanorbis*. One postulates that a part of the ensemble of species has a Gondwanan vicariant origin (Meier-Brook, 1984). The other hypothesis emphasizes on dispersal of *proto-Indoplanorbis* into India from Africa through West Asia (Arabian Peninsula) or Central Asia through Sinai-Levant tract after the collision of Africa with Eurasia (Gauffre-autelin et al., 2017; Liu et al., 2010; Morgan et al., 2002). Molecular phylogenetic studies in conjunction with divergence dating methods and an absence of Cretaceous Bulinid fossils from India hint towards the latter hypothesis being more plausible (Gauffre-autelin et al., 2017). Whereas, Liu et al., (2010) inferred a Miocene-middle Pleistocene dates for diversification for the *I. exustus* clades, a later study estimated that the diversification took place earlier during mid-Miocene to early Pleistocene (Gauffre-autelin et al., 2017). Furthermore, studies by Gauffre-autelin et al., (2017) also suggested presence of five cryptic lineages present in the species complex as opposed to two as suggested by Liu et al., (2010). Interestingly, all these studies concluded that basal diversification took place in the humid plains of Nepal or North India (Gauffre-autelin et al., 2017; Liu et al., 2010). It was also inferred that late Miocene intense aridification, which occurred owing to upliftment of the Quinghai-Tibetan Plateau (QTP) and orogenesis in the Himalayas, and the Quaternary glaciation induced drying is responsible for many of these speciation events (Gauffre-autelin et al., 2017; Liu et al., 2010). These authors also pointed out that major river and lake systems might have served a role in shaping their current distribution pattern. (Gauffre-autelin et al., 2017), noted that clade five from their study, which is widespread in Asia, underwent extensive population expansion in the recent past. This study also found no population genetic structure in clade five from the Sundaic Islands, suggesting that the species can easily disperse across known biogeographic barriers.

These studies provided a broad overview, however, a thorough sampling of the Indian subcontinent, where the species complex is widespread and potentially harbours much genetic diversity, is necessary. Furthermore, during the Pleistocene, xerification of a large part of North-western India led to fragmentation of forest and freshwater habitats (Roberts et al., 2018). It is likely that these climatic fluctuations in the subcontinent shaped the population history of the members of the species complex after their divergence from each other. The complex topography of the subcontinent also creates a range of geographic barriers which might generate population genetic structure. A greater representation of populations from India might provide a better understanding of the number of putative species, the centre of diversification of the species complex, the causal factors driving the radiation, as well as the population genetic history of each species after their divergence from the sister species. To address some of these issues we sampled *Indoplanorbis* populations from all the major river basins of India to better understand their origin and evolution. We carried out species delimitation, fossil calibrated molecular dating analysis, and ancestral range estimation analyses, to conclusively estimate the historical biogeography of the species complex. Furthermore, we also implement population genetic and statistical phylogenetic methods. The goal of this study was to address the following questions – 1) When did *Indoplanorbis* colonize India and from where? 2) How many putative species constitute the species complex? When and where did basal radiation in the complex begin? 3) What are the climatic and geographic factors that contributed to the radiation event? 4) How did *Indoplanorbis* disperse across Asia after the initial radiation, 5) Did any of the putative species undergo population contraction and subsequent expansion during the Pleistocene glacial cycles? 6) Is there any population structure observed in the species complex as a whole or in any of the putative species?

## Materials and Methods

### Sample Collection

*Indoplanorbis* samples were collected from a diverse range of lotic and lentic habitats. Samples were collected from Indian subregion and Northeast India which is part of Indo-Chinese subregion. Furthermore, Multiple river basins were sampled from the Indian subregion (see Appendix B for a complete list of samples). The latitude and longitudes were noted while sample collection. The samples were preserved in absolute alcohol for further analysis.

### Molecular work

Genomic DNA was extracted from the preserved samples using CTAB extraction method (Chakraborty et al. 2020) and thereafter quantified using nanodrop. A partial fragment of COI gene was PCR amplified from all the extracts. Furthermore, the H3 and 18S genes were amplified only from one individual from each putative species (see Table 1 for a complete list of molecular markers used). Each 25 μL reaction contained ~60 ng template DNA, 0.3 μM of each primer, 0.25 μM dNTPs mixture, 0.04 μg/mL BSA, 2.5 μM MgCl2, 1X Taq buffer, and 1.5U Taq DNA polymerase. The rest of the volume was made up of Mq water. The PCR steps performed are as follows — 3 minutes at 95°C, 45 seconds at 94°C, 45 seconds at 41–54°C, 2 minutes at 72°C, 10 minutes at 72°C, and finally 4°C for an indefinite period of times. Step 2 to 5 were repeated 39 times. The annealing temperature at step 3 varied between different primers (see Table 1 for details) Agarose gel electrophoresis was used to observe the successful amplification of the DNA. Thereafter, the PCR products were sent to Barcode Biosciences, Bangalore for sanger sequencing.

**Table 1.** List of genes.

| Gene name | Abbreviation | Length (bp) | Models | Primers | References |
| --- | --- | --- | --- | --- | --- |
| Cytochrome c oxidase subunit 1 | COI | 589 | Codon position 1 HKY, codon position 2 HKY+I+G, codon position 3 TrNef+I | LCO and HCO | (Folmer et al., 1994; Williams et al., 2003) |
| 18S ribosomal RNA | 18S rRNA | 366 | TrNef+I | 18SYLMFOR<br>18SYLMREV | (Stothard et al., 2000) |
| Histone H3 | Histone H3 | 303 | Codon position 1 and 2 TrNef+I, codon position 3 K80+I | H3F and H3R | (Colgan et al., 1998) |

### Phylogenetic analysis

The quality of the sequence reads obtained were checked on Chromas 2.6.6 (Technelysium Pty Ltd.). Next, the sequences were imported to MEGA7 (Kumar et al., 2016) and aligned using MUSCLE (Edgar, 2004) in combination with *Indoplanorbis exustus* sequences from all over its distribution range and *Bulinus* sequences obtained from online repositories. Several species of *Bulinus* served as outgroup. The models of sequence evolution and the correct partition scheme were estimated in PartitionFinder2 (Lanfear et al., 2017). The maximum likelihood (ML) analysis was carried out using RAxML HPC 1.8.2 (Stamatakis, 2014) implemented in raxmlGUI 1.5 (Silvestro and Michalak, 2011). Next, a single locus based species delimitation method, multi-rate Poisson Tree Process (mPTP) (Kapli et al., 2017) was employed to estimate the number of putative species in the species complex, based on the ML tree. A concatenated dataset was built from one individual from each putative species in Mega7 for downstream analyses.

### Divergence dating

Divergence dating was carried out in order to uncover the biogeographic and lineage history of the species complex. The analyses were carried out in BEAST 1.8.3 (Drummond et al., 2012). The analyses were based on the concatenated dataset. A total of two Miocene fossil calibrations were used: the oldest fossil from *Bulinus africanus* group (~19 million years ago or mya) and the oldest fossil from *Bulinus tropicus* group (~15.5 mya). We assumed the MRCA (most recent common ancestor) from each group would be older than the age of the fossils, hence, gamma distribution priors were specified for each calibration. The fossils only convey information about the minimum bound of the calibration, not about the mode of the distribution. Hence, to incorporate the uncertainty about the mode of the distribution, we employed a combination of shape (α) parameter values in four independent but otherwise identical runs and compared the marginal likelihood of each run using Bayes factor test. The combinations are as follows: A. α parameter value of 2 for the MRCA of both *B. africanus* and *B. tropicus* groups. 2. α parameter value of 2 for the MRCA of *B. africanus* group and 4 for the MRCA of B. *tropicus* group. 3. α parameter value of 4 for the MRCA of *B. africanus* group and 2 for the MRCA of *B. tropicus* group. 4. α parameter value of 4 for the MRCA of both *B. africanus* and *B. tropicus* groups. The scale (β) parameter value was fixed at 2 for all runs. The marginal likelihood estimates were computed using path sampling and stepping stone sampling method. A hundred path steps were performed for 5,00,000 generations. The saturation of each BEAST run was visually checked in Tracer v1.7.1. Furthermore, ess values >200 was considered as decent sampling of the parameter space. The trees were summarized in TreeAnnotator 1.8.0 and the first 25% of the trees were discarded as burnin. Finally, the maximum clade credibility tree was visualized in fig Tree.

### Ancestral area reconstruction analysis

Ancestral range estimation analyses were carried out using the package BioGeoBEARS 1.1 in R 3.4.2 (available at https://www.r-project.org/) based on the maximum clade credibility tree from the divergence dating analysis with the highest MLE. We implemented the DEC+*j* models to assess the range inheritance patterns. The model has performed well for freshwater snail biogeographic reconstruction previously (Sil et al., 2020). The DEC+*j* model is particularly useful while estimating the ancestral ranges of dispersal limited taxa and island taxa (Hendriks et al., 2019; Kitson et al., 2018). Some of the landmasses where Bulinidae is distributed in, were isolated island landmasses during part of the group’s biogeographic history. Furthermore, freshwater snails also exhibit avian-mediated dispersal which can be best modelled by the founder effect speciation incorporated in the DEC+*j* model. The entire distribution range of *Indoplanorbis* and *Bulinus* were partitioned into three regions that roughly correspond to Biogeographic regions and subregions, namely, Southeast Asia (Sundaland, Mainland Southeast Asia, Northeast India), India (Sri Lanka, Nepal and India apart from Northeast India), Africa. Direct dispersal from Africa to Southeast Asia and vice versa were assigned a low dispersal probability, since these landmasses were not adjacent during the time period taken into consideration here (40–0 mya). Furthermore, we divided the time period into two time slices following Sidharthan and Karanth (2021). These are 40–35 mya, when Africa and India did not have permanent land connection with Eurasia (Aitchison et al., 2008; Allen and Armstrong, 2008), and thus an ancestral range consisting of any two of the biogeographic areas would not have been possible; 35–0 mya, after the collision of Africa and India with Eurasia, when an ancestral area consisting of a combination of geographic areas will be possible (see Table A1 for details of the BioGeoBEARS analyses).

### Estimation of population size change

The historical population size fluctuations in each putative species were estimated in order to find out whether the climatic fluctuations in the Quaternary period affected any of the putative species. Species 1 and 3 were excluded owing to small sample size. We know from previous literature (Gauffre-autelin et al., 2017), that crown radiation in these species began during Pleistocene. Hence, it is reasonable to assume that the population size fluctuations in each putative species would result from the climatic changes during the Pleistocene. Different summary statistics were calculated in Arlequin ver 3.5.2.2 (Excoffier and Lischer, 2010). Furthermore, mismatch distribution was implemented in Arlequin. Finally, population size change over time was estimated using Bayesian skyline plot implemented in BEAST 2.6. We used a substitution rate of 1.57 % per million years as clock rate following Wilke et al., (2009). The birth death skyline contemporary model was employed since we were sampling from multiple population containing homochromous data (when all the individuals were sampled during the same time period). The birth death model is more powerful in handling large sampling sizes than the coalescent model (Stadler et al., 2013). A beta distribution was selected for the rho prior. The values of alpha and beta were set at 1 and 9999 respectively, as our sampled individuals were a small fraction of the total population. A lognormal distribution was employed for the origin prior. The mean was set to 14.5 and standard deviation to 0.5 in order to move the median of the distribution around 2 mya. Each analysis was carried out for 10 million generations. The saturation of each run was visually checked at Tracer and from ESS values. Finally, the Bayesian skyline plots were visualized in R using the package bdskytools.

### Estimation of genetic structure

We aimed to understand the influence of climatic and geographical barriers on the population structure of *Indoplanorbis*. The population structure of each putative species was estimated in BAPS6 (Corander et al., 2004) in a Bayesian framework. Species 1 and 3 were excluded owing to small sample size. The ‘clustering of linked loci’ algorithm was implemented to estimate the population structure. Since, we did not have any prior information about the plausible number of clusters, each analysis was repeated, setting the number of clusters to five and ten respectively.

## Results

### Phylogenetic and species delimitation analyses

The ML reconstruction obtained a well-supported phylogeny of *Indoplanorbis* consisting of six clades (Figure 1). The phylogeny retrieved two major clades, one confined to Indian subcontinent and the other in both Indian subcontinent and Southeast Asia. The Indian clade consists of two subclades. Clade one was restricted only to Northern India, whereas clade 2 was distributed in Northern and Peninsular India as well as Nepal. Amongst the members of the other major clade, Clade 3 consisted of individuals from Laos. Clade 4 was widespread in the subcontinent from Bangladesh, Odisha, northeast Indian states to Nepal. Clade 5 was widespread in Southeast Asia including Indo-China, Malayan Peninsula and Sundaland. It was also distributed in Peninsular India, Nepal and Oman in West Asia. Clade 6 was restricted to Myanmar in Southeast Asia, parts of India and Nepal. The results from the mPTP species delimitation analysis suggest six putative species, each corresponding to the six clades described above. Henceforth, each clade will be referred to as putative species bearing the same number.

**Figure 1:**
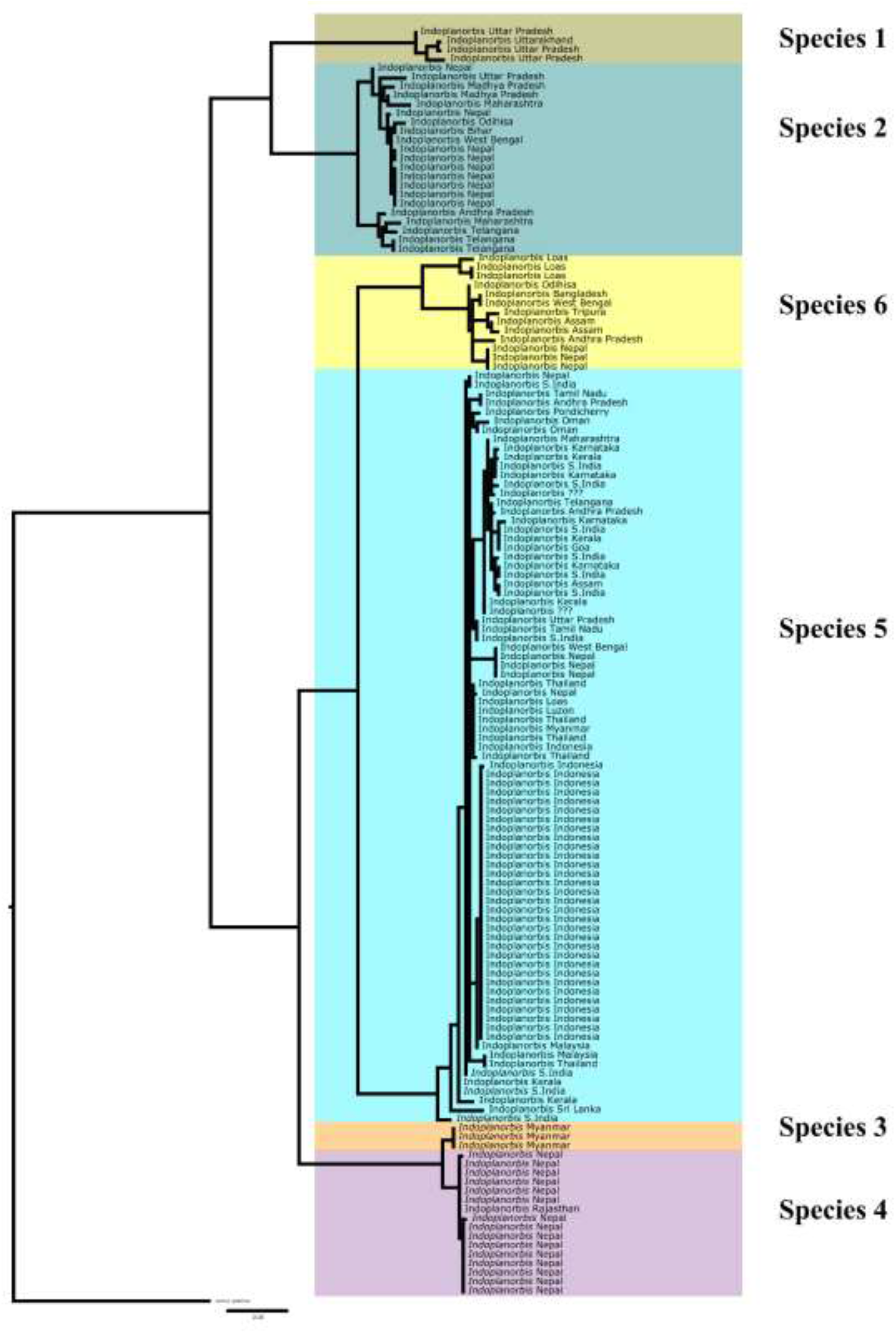
ML tree showing the six putative species that constitute *Indoplanorbis exustus species* complex

### Divergence dating

The divergence dates obtained from analyses aided by fossil calibrations were fairly consistent with the results from previous studies based on substitution rate (Figure 2). Out of the four analyses, the one guided by the third model (α parameter value of 4 for the MRCA of *B. africanus* group and 2 for the MRCA of *B. tropicus* group) had the highest MLE, however, it was not strongly supported in the Bayes Factor test (see Table A2). Since, this model had the highest log likelihood, henceforth, we will discuss the results from the third model. *Indoplanorbis* diverged from *Bulinus* during Eocene–Oligocene (51.9–27.6 mya). Basal radiation in the *Indoplanorbis* complex began during Oligocene–Miocene (31.4–14.07 mya). Putative species A diverged from B during Miocene (21.2–7.5 mya). Putative species 6 split off from the MRCA of 5, 4 and 3 around the same time (21.6–9.1 mya). Later on, putative species 5 separated from the MRCA of 3 and 4 during mid to late Miocene (15.3–6.0 mya). The most recent divergence took place between putative species 3 and 4 around late Miocene– Pleistocene (7.3–2.2 mya).

**Figure 2:**
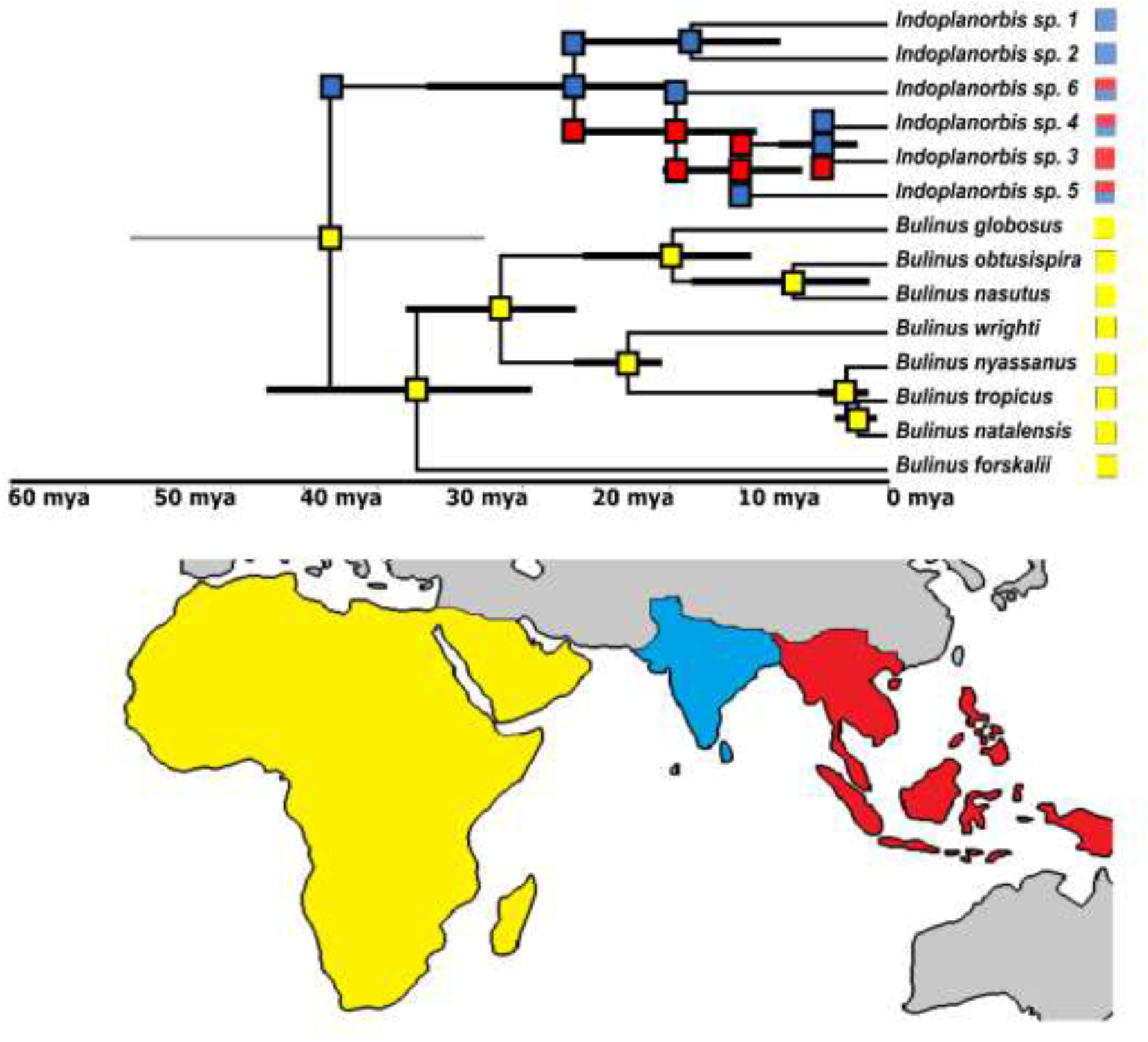
A chronogram showing the historical range evolution of *Indoplanorbis*

### Ancestral area reconstruction analyses

According to the BioGeoBears analysis, *Indoplanorbis* lineage dispersed into Asia from Africa sometime during Eocene–Oligocene (51.9–27.6 mya) (Figure 2). The initial radiation took place in India during Oligocene–Miocene (31.4–14.07 mya). The lineage consisting of putative species 3, 4, 5 and 6 dispersed into Southeast Asia around the same time. Several back dispersals to India took place after that starting from mid-Miocene till Pleistocene.

### Estimation of population size change

the population size estimation analyses under various methods yielded mixed results (see Table A3). Tajima’s D and Fu’s Fs were negative for all four putative species analysed. Whereas Fu’s Fs were significant for all the species but *Indoplanorbis* species 4, only *Indoplanorbis* species 5 came out as significant in Tajima’s D. The Bayesian skyline plots showed some degree of population fluctuation in the demographic histories of all putative species (Figure A3). The mismatch distribution graphs of all species were multimodal suggesting that the populations have been stable over time (Figure A2). However, the squared sum of standard deviation (SSD) values of the mismatch distribution calculated under the demographic expansion model were not significantly different in case of any species which suggests recent demographic expansion.

### Estimation of genetic structure

All the species exhibited population breaks, however, they could not be linked to any known biogeographic barriers (Figure A1). Individuals from *Indoplanorbis* species 2 formed three clusters. Individuals from the first two cluster consisted of individuals from Peninsular India. The third cluster also consisted of individuals from Peninsular India as well as Gangetic basin of India and Nepal. *Indoplanorbis* species 4 had four clusters. The first cluster consisted of individuals from Nepal. The second clusters consisted of individuals mainly from Northeast India and Bangladesh. The third cluster consisted of individuals from Northeast India, eastern Gangetic basin, as well as Peninsular India, whereas the fourth cluster consisted of individuals only from eastern Peninsular India. *Indoplanorbis* species 5 was composed of three clusters. The first cluster consisted exclusively of individuals from Peninsular India, and the third cluster solely of individuals from mainland Southeast Asia and Sundaland. The second cluster included individuals from all these regions. *Indoplanorbis* species 6 consisted of three clusters. The first three clusters consisted of individuals from Rajasthan and Nepal respectively, whereas the fourth cluster consisted of individuals from Nepal and Myanmar.

## Discussion

In this paper we implement several biogeographic, phylogeographic and population genetic tools to trace the lineage history of *Indoplanorbis*. We found out that *Indoplanorbis* is a species complex consisting of six putative species, which colonized the Indian subcontinent during Eocene–Oligocene. The results show that basal radiation in the species started during Late Oligocene–Miocene in India. We further demonstrate that the putative species exhibit little genetic structure, however, all the species underwent population fluctuations during the last 2 million years. Below, we discuss the possible factors governing the patterns we uncovered.

### Colonization of India

Several hypotheses were put forward to explain the time frame and route of colonization of *Indoplanorbis* into India (Attwood et al., 2007; Gauffre-autelin et al., 2017; Liu et al., 2010). The Gondwanaland origin theory were not supported by any of the studies published before. Here, for the first time, we used a fossil calibrated phylogeny and statistical methods to demonstrate that colonization dates definitely postdate the continental separation between Africa and India, thus refuting Gondwanaland origin hypothesis (Figure 2). Although, all the studies have agreed upon a dispersal timeframe which took place after the collision of Africa and India with the Eurasian plate, the actual timeframe widely varies. Previous studies have suggested a Eocene-Miocene dispersal or a Pleistocene dispersal from Africa via the Sinai-Levant dispersal tract (Attwood et al., 2007) or through Arabian Peninsula (Morgan et al., 2002). Our results showed an Eocene-Oligocene dispersal which corroborates the previous findings (Attwood et al., 2007; Gauffre-autelin et al., 2017; Morgan et al., 2002). The dispersal pattern observed here also closely resembles the dispersal of strepsirhines (Yoder and Yang, 2004) and *Pila* (Sil et al., 2020) from Africa to tropical Asia. It is likely that *Indoplanorbis* dispersed during the Eocene-Oligocene boundary, which is marked by the beginning of the closing of the peri-Tethys ocean and collision of Afro-Arabian plates with Eurasia (Meulenkamp and Sissingh, 2003). The climate in Central and West Asia was also more conducive to the dispersal of freshwater species around that time as evident from the fossil record and paleoclimatic reconstructions (Stothard et al., 2000) Harzhauser et al., 2016; Kraatz and Geisler, 2010; Neubert and Damme, 2012).

### Diversity of *I. exustus*

Previous studies hinted at *Indoplanorbis exustus* being a species complex (Devkota et al., 2015; Gauffre-autelin et al., 2017; Liu et al., 2010). It is also generally observed that widespread species commonly harbour cryptic diversity (Karanth, 2017). *Indoplanorbis*, which is distributed from Iran to China, would be prime candidate of such examination. Liu et al. (2010) described three lineages of *Indoplanorbis*, although he was sceptical about designating species status to all of them. Later Devkota et al., (2015) added more samples chiefly from Nepal and demonstrated the presence of four lineages in *Indoplanorbis* species group. Gauffre-autelin et al. (2017) included a much wider range of samples and showed that *Indoplanorbis* consists of five lineages. Both the studies suggested that *Indoplanorbis* is perhaps a species complex consisting of multiple putative species. Here we showed the presence of six putative species on the basis of molecular species delimitation method (Figure 1). However, more lines of evidence have to be used to fortify this conclusion. Morphology based classification has failed previously to characterise putative species in this species complex (Devkota et al., 2015). Anatomy, cytogenetics and niche modelling techniques need to be implemented in order to estimate the actual diversity in this species complex.

Northeast India, Northern India or Nepal was suggested as regions where the initial phase of diversification in *Indoplanorbis* took place by previous studies (Gauffre-autelin et al., 2017; Liu et al., 2010). The ancestral area reconstruction analyses suggest that MRCA of the complex was distributed in India. The distribution range of all the putative species included parts of the subcontinent. Moreover, *Indoplanorbis* species 1 and 2 which formed one of the two major clades in the complex were exclusively found in the subcontinent. Putting together all these, it is likely that the diversification started somewhere in the Indo-Gangetic plains of Northern India or Nepal.

There have been many hypotheses concerning the drivers of diversification in this group. Liu et al., (2010) estimated the beginning of diversification in this group to be ~7 mya. The study attributed to Himalayan upliftment (~8 mya) which caused strengthening of the Asian monsoon and aridifcation during the late Miocene as the driver of the initial divergence in the group. This study also suggested that *Indoplanorbis* requires humid climate with plenty of waterbodies to sustain a viable population. The aridifcation might have led to population contraction into multiple refugia and subsequent allopatric speciation. The more recent divergences were attributed to mid-Pleistocene climatic fluctuations and resultant aridification events (Liu et al., 2010). Gauffre-autelin (2017) obtained much earlier divergence dates (~16 mya) from a much bigger dataset. The study by Gauffre-autelin et al. (2017) suggested that the intensification of Asian monsoon periodically from mid-Miocene to Pliocene about 15–13 mya, 8 mya and 3 mya, was the primary driver of divergence in this group. These intensification events resulted from the Himalayan orogeny, the retreat of the Paratethys sea from Central Asia, and the onset of the first major glaciation event in the Northern Hemisphere during Pliocene (Tada et al., 2016; Zhang et al., 2009). These events roughly coincide with the three initial divergence dates obtained by Gauffre-autelin (2017). The final divergence was attributed to the aridification resulting from Quaternary climatic oscillations (Gauffre-autelin et al., 2017). The dates obtained from the current study are congruent with those obtained from Gauffre-autelin et al. (2017). Hence, it seems likely that the Asian monsoon driven aridification events might have been responsible for driving allopatric speciation in this species complex. The population genetic analyses carried out in the current study further corroborate this conjecture. All the putative species showed signatures of population fluctuation during the last 2 my, which suggests these organisms are susceptible to climatic oscillations. Hence, it is highly likely that even older events might have affected the stem lineages in similar fashion and brought about the speciation events. Aridification is a known driver of allopatric separation in freshwater organisms (Beheregaray et al., 2017; Buckley et al., n.d.). In the Indian subcontinent aridification-driven allopatric speciation was documented in Ichthyophid caecilians (Gower et al., 2016). Casual observation shows *Indoplanorbis* can be found in a wide range of habitat, including ephemeral waterbodies suggesting they would be less susceptible to desiccation than other freshwater organisms. However, a few experimental studies showed that the survival of *Indoplanorbis* is affected by low temperature (Raut et al., 1992). Low temperature is typical of periods of increased aridity (Zachos et al., 2001). Moreover, juveniles of *Indoplanorbis* exhibit high mortality in response to desiccation (Parashar and Rao, 1982). Hence, it is plausible that despite being a generalist species, *Indoplanorbis* underwent range contraction and allopatric separation during mid-Miocene to Pleistocene. Gauffre-Autelin et al., (2017), further suggested that geographical features such as Bramhaputra and Mekong river basin might have played a role in partitioning genetic diversity and subsequent speciation. The current study, however, could not detect any genetic structure in any of the putative species. *Indoplanorbis* is a pulmonate snail which can breathe air and appear to be capable of tiding over unfavourable conditions. This ability makes it a more potent disperser over-land. This would also aid in successful bird-mediated dispersal events. All the *Indoplanorbis* species seem to be quite generalists occupying a large range of habitats. River basins might be a more effective barrier to dispersal for less vagile species.

### Effects of Pleistocene climatic cycles and geographic barriers

Phylogeography of flora and fauna in the temperate regions were largely shaped by the Pleistocene climatic oscillations (Avise, 2000; Hewitt, 2004). However, the effects of the climatic oscillations manifested in the tropics differently. The glaciation events in higher latitudes led to cycles of aridification and lowering of sea level in the tropical and subtropical regions. This in turn resulted in population expansion or contraction, dispersal of terrestrial taxa to newly connected landmasses, and population divergence in marine species. For example, land connections were established between various Sundaic Islands and mainland Southeast Asia, as well as between Sri Lanka and India several times during the Pleistocene which resulted from the aforementioned lowering of the sea level (Bossuyt, 2014; Brown et al., 2013). In the Indian subcontinent many taxa which favours humid climate underwent range contraction and local extinction during the glacial phases (Iyengar et al., 2005; Natesh et al., 2020; Vidya, 2016). Other lines of evidence from palynology and sedimentation studies as well as paleoclimate modelling also suggest xerification in many parts of the subcontinent especially Northwestern India (Goodbred, 2003; Roberts et al., 2018). Previously Gauffre-autelin et al., (2017) showed that the Sundaic clade of putative species 5 underwent population expansion after a bottleneck which coincided with the Pleistocene glacial events. Here we investigate the effects of the glacial driven aridification on each of the species except *Indoplanorbis* species 1 and 3 owing to small sampling size. The Fu’s Fs were negative and significant for all species except *Indoplanorbis* species 4, whereas Tajima’s D was negative and significant for only *Indoplanorbis* species 5 (Table A3). The mismatch distribution analyses showed non-significant P values of SSD and raggedness index under the demographic expansion model suggesting that sudden population expansion has taken place in all four species of *Indoplanorbis* (Table A3 and Figure A2). Similarly, the Bayesian skyline plots (Figure A3) showed population fluctuations in the history of all putative species although at different time points. To conclude, all the species were likely to have underwent population crash and subsequent expansion during Pleistocene. The most likely cause must have been the aridification event in South and Southeast Asia. However, much finer scale sampling and broader gene coverage is necessary to understand, in which parts of the range of each species, population contraction took place and which regions served as refugia. Such patterns are expected from species such as *Indoplanorbis* spp. which is dependent on freshwater sources which were likely to have depleted during the aridification cycles. A lot of freshwater species groups exhibit such patterns in response to the Pleistocene glaciation driven aridification events (Bagley et al., 2013; Buckley et al., n.d.; Guzik et al., 2012; Klunzinger Manuel Lopes-Lima Andre Gomes-dos-Santos Elsa Froufe A J Lymbery L Kirkendale, n.d.; Unmack et al., 2012; Ye et al., 2018). However, our field observation suggested that *Indoplanorbis* can survive in ephemeral bodies of water as well. Hence, experimental studies need to be carried out in order to ascertain to what degree *Indoplanorbis* can withstand desiccation of its habitat. A handful of experimental studies suggest that *Indoplanorbis* is sensitive to desiccation and low temperature often associated with aridification events.

Interestingly, none of the species exhibited any obvious spatial genetic structure (Figure A1). Their ability to breathe air might have enabled them to disperse across river basins and other biogeographic barriers. Moreover, their small size might have enabled them to be dispersed successfully by birds after attaching to their feathers and legs or after getting ingested. However, it remains to be seen if any other aspect of their biology such as physiological tolerance or life history contributed to their successful dispersal and colonization.

### Conclusion

In this paper we showed that *Indoplanorbis* is most likely not a single species but a species complex consisting of six putative species, as suggested by the molecular species delimitation method. The lineage leading to *Indoplanorbis* colonized India from Africa during Eocene–Oligocene, most likely during late Eocene or early Oligocene, as that time the geological arrangement of landmasses and the climatic conditions were conducive to the dispersal of freshwater species. The lineage dispersed through Central or West Asia following the Eurasian route similar to *Pila*. The basal diversification took place in India during late Oligocene–Miocene. This and the subsequent divergence events in the lineage were catalysed by aridification events caused by the strengthening of the Indian monsoon and latter Quaternary aridification cycles driven by global changes. Each putative species exhibits signatures of rapid population growth which is most likely to have been caused by the Pleistocene glaciation cycles driven aridification events. We did not observe any explicit population genetic structure suggestive of a highly vagile species. A finer scale sampling and greater gene sampling strategy might uncover the probable population genetic structure and the location of the Pleistocene refugia occupied by each species.

## Acknowledgement

The major part of the project was funded by grants from Dept. of Biotechnology, Govt. of India (BT/01/17/NE/TAX) to NAA. The part of the fieldwork was supported by Rufford Small Grant for Nature Conservation (19805-1) to MS and DST (Department of Science and Technology, Govt. of India)-SERB grant (EMR/2017/001213) to PK. Authors would like to thank Pruthviraj, Harshal Bhosale, Tarun Singh, Aparna Lajmi, Chinta Sidhharthan, Kunal Arekar, Ananya Jana, Surya Narayan, Suhel Seikh, Animesh Mondal and Debottam Bhattacharjee for help during sample collection. Chinta Sidhharthan provided important inputs towards BioGeoBEARS analysis.

