## Appendix A for "Historical biogeography and phylogeography of *Indoplanorbis exustus*"

**Table A1.** Details of BioGeoBEARS analyses whowing the adjacency and dispersal probability between different landmasses at different time frames

| Time frames (mya) | Continents | Adjacency | Areas Allowed | Dispersal probability | Relative position of continents | References |
|---|---|---|---|---|---|---|
| 52–35 | India-Africa | 1 | 0 | 1 | Before the permanent land connection was established between Africa and Eurasia, as well as India and Eurasia; biotic interchange was possible between the landmasses through temporary landbridges or jump dispersal. However, an ancestral area consisting of two or more landmasses was not possible as the landmasses were not permanently connected. | (Aitchison et al., 2008; Allen and Armstrong, 2008) |
| | India-SEA | 1 | 0 | 1 | | |
| | Africa-SEA | 0 | 0 | 0.01 | | |
| 35–0 | India-Africa | 1 | 1 | 1 | Permanent land connection was established between Africa and Eurasia, as well as India and Eurasia. Biotic interchange was possible between all the landmasses. An ancestral area consisting of India-Africa and India-SEA was possible since, there was permanent land connection between these landmasses. | |
| | India-SEA | 1 | 1 | 1 | | |
| | Africa-SEA | 0 | 0 | 0.01 | | |

**Table A2.** Bayes Factor test (*B. africanus* group is abbreviated as A, *B. tropicus* group as T; the numbers next to the names denote the value of the a parameter of the gamma distribution; the numbers within brackets are the marginal likelihood estimates; the numbers below are the log Bayes Factor values)

| MLE from path likelihood | | | | |
|---|---|---|---|---|
| | A4T2 (-3901.8) | A4T4 (-3902.09) | A2T2 (-3902.43) | A2T4 (-3904.03) |
| A4T2 | | | | |
| A4T4 | 0.29 | | | |
| A2T2 | 0.63 | 0.34 | | |
| A2T4 | **2.23** | **1.94** | **1.6** | |
| MLE from stepping stone | | | | |
| | A4T2 (-3901.86) | A4T4 (-3902.2) | A2T2 (-3902.44) | A2T4 (-3904.02) |
| A4T2 | | | | |
| A4T4 | 0.34 | | | |
| A2T2 | 0.58 | 0.24 | | |
| A2T4 | **2.16** | **1.82** | **1.76** | |

**Table A3.** Population genetic summary Statistics

| | Tajima's D | P value | Fu's Fs | P value | SSD | P value | Raggedness index | P value |
|---|---|---|---|---|---|---|---|---|
| Species 2 | -1.456 | 0.054 | -6.87 | **0.008** | 0.0125 | 0.7 | 0.011 | 0.87 |
| Species 4 | -0.62 | 0.276 | -1.21 | 0.153 | 0.029 | 0.74 | 0.027 | 0.92 |
| Species 5 | -1.356 | **0.048** | -24.008 | **0.001** | 0.019 | 0.25 | 0.033 | 0.13 |
| Species 6 | 0.197 | 0.47 | -9.32 | **0.001** | 0.072 | 0.34 | 0.127 | 0.46 |

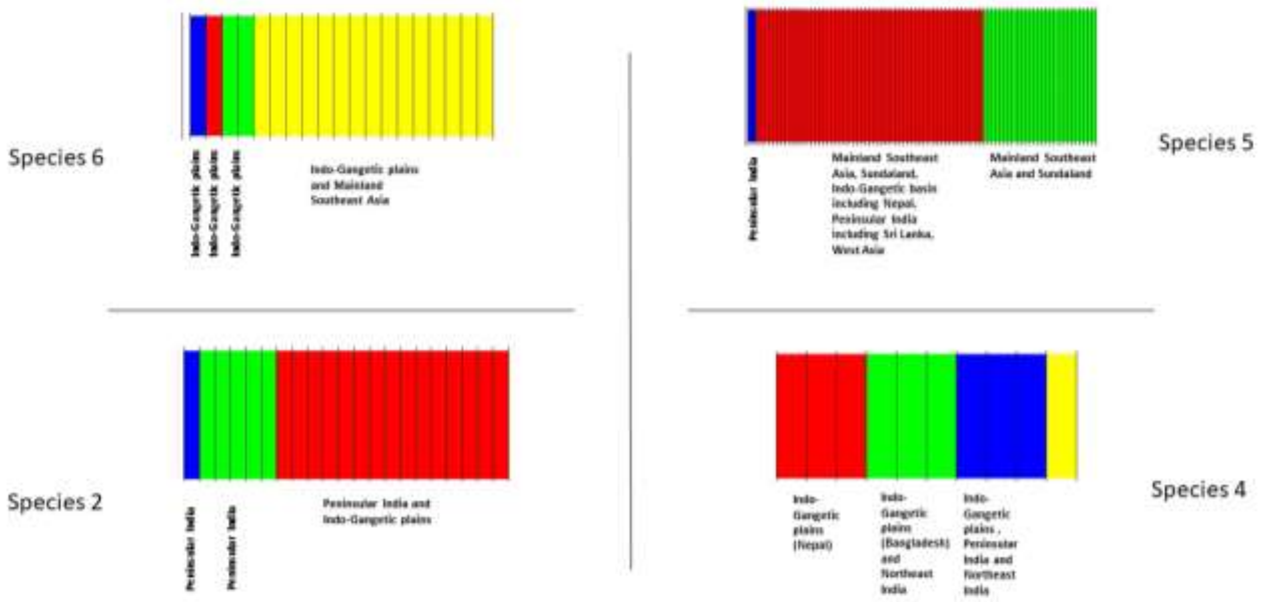


**Figure A1:** Genetic structure in each putative species as estimated by BAPS6. Each coloured section in the figure represents a genetic cluster.

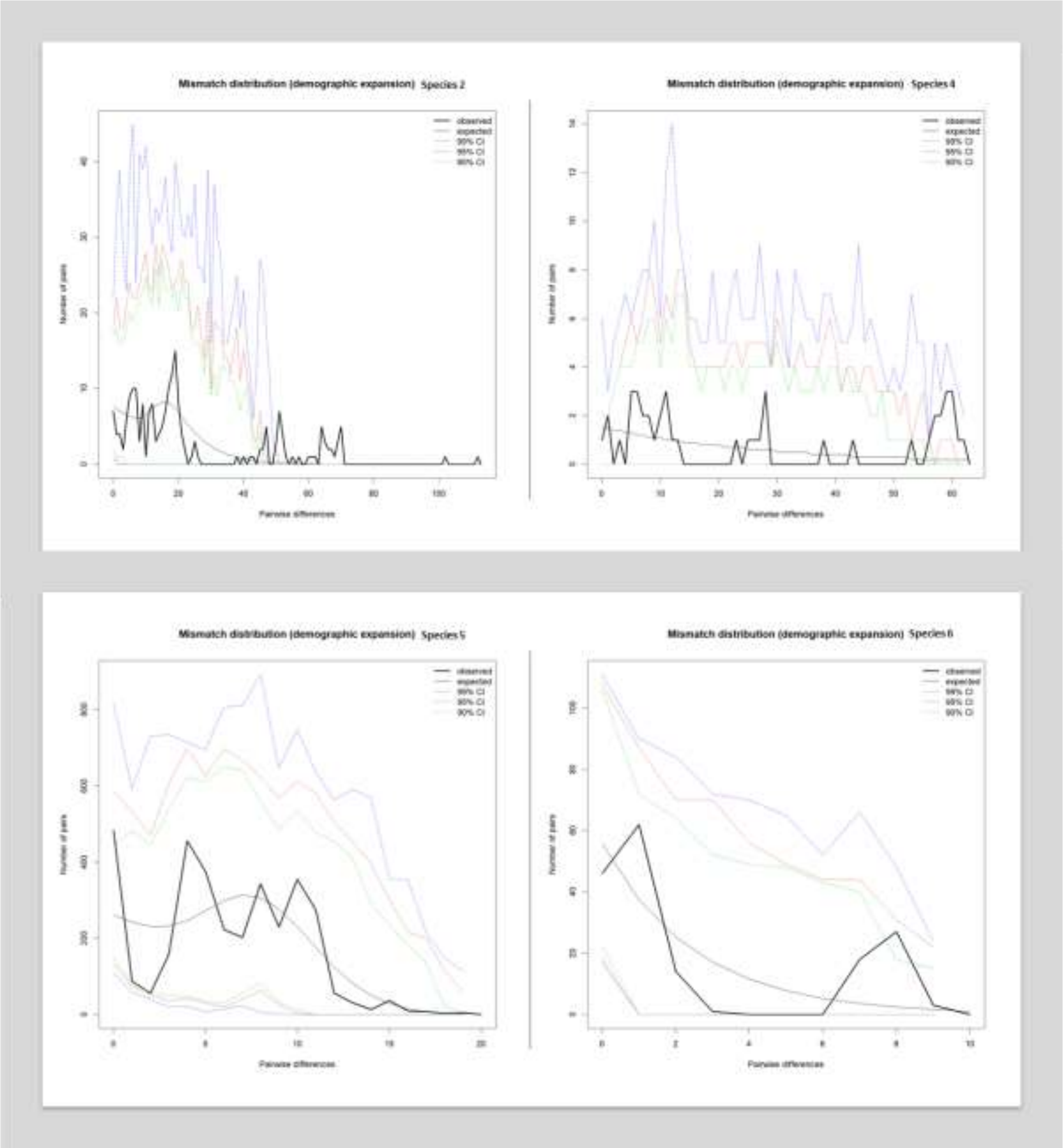


**Figure A2:** Mismatch distribution plot under the assumption of demographic expansion for each putative species. The solid line represents the observed distribution

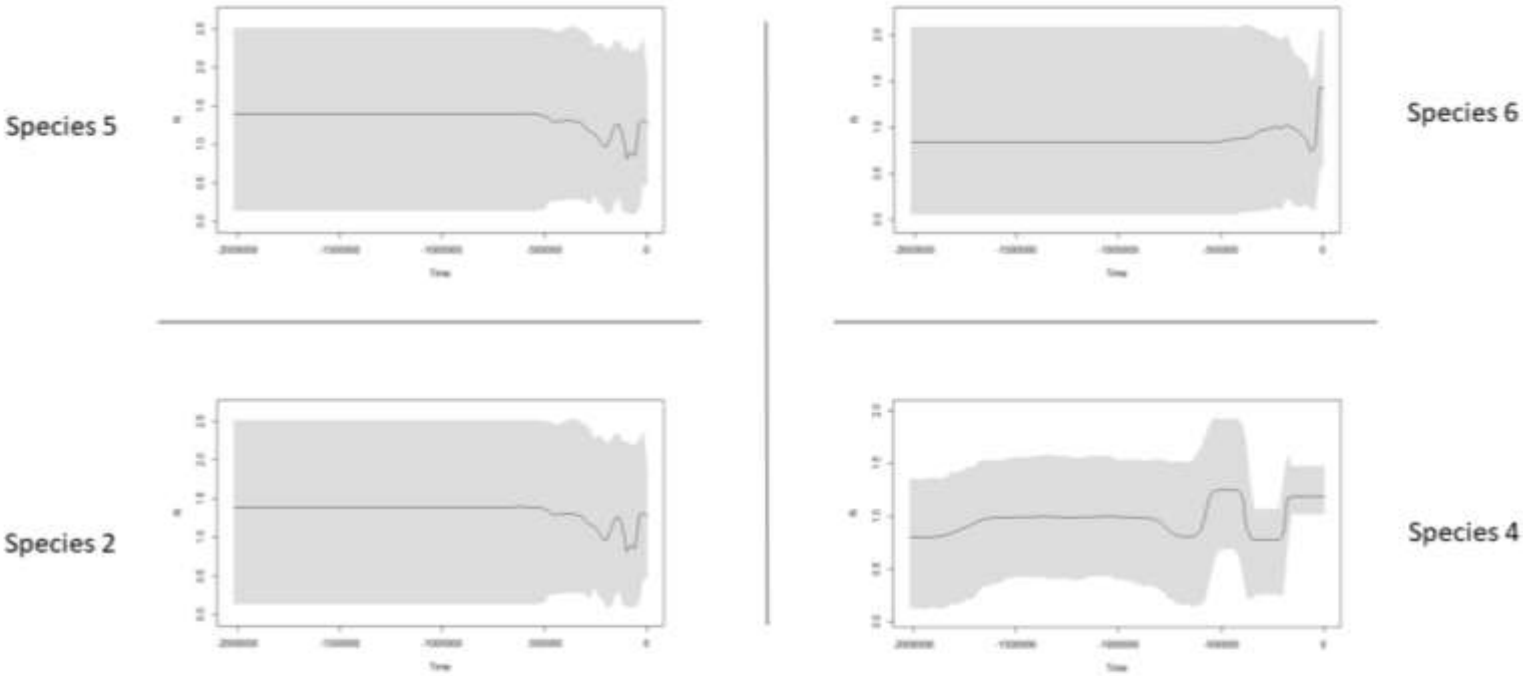


**Figure A3:** Bayesian Skyline plots of each putative species.
