## Appendix B for "Historical biogeography and phylogeography of *Indoplanorbis exustus*"

**Table B1: Sequences generated during the current study**

| S.No | SAMPLE ID | Country | SAMPLE LOCATION |
|---|---|---|---|
| 1 | MS 452.1 | India | Coorg, Karnataka |
| 2 | MS 566.1 | India | Pondicherry, Tamil Nadu |
| 3 | MS 489.1 | India | TVM, Kerala |
| 4 | MS 428.1 | India | Panaji, Goa |
| 5 | MS 465 | India | Palakkad, Kerala |
| 6 | MS 491 | India | Kalpetta, Kerala |
| 7 | MS 236 | India | Bhubaneswar, Odhisa |
| 8 | MS 543 | India | Pedapalli, Telangana |
| 9 | MS310 | India | Wayanad, Kerala |
| 10 | MS553 | India | Kurnool, Andhra Pradesh |
| 11 | MS 292 | India | Guna, Madhya Pradesh |
| 12 | MS 545 | India | Hyderabad, Telangana |
| 13 | MS 522 | India | Khammam, Telangana |
| 14 | MS 497 | India | Ongole, Andhra Pradesh |
| 15 | MS 531 | India | Mancherial, Telangana |
| 16 | MS 334 | India | Nagpur, Maharashtra |
| 17 | MS 403 | India | Halekote, Karnataka |
| 18 | MS 472 | India | Palakkad, Kerala |
| 19 | MS 605 | India | Roorkee, Uttarakhand |
| 20 | MS 625 | India | Raiganj, West Bengal |
| 21 | MS 158 | India | Sawai Madhopur, Rajasthan |
| 22 | MS 110 | India | Mirzapur, Uttar Pradesh |
| 23 | MS 94 | India | Patna, Bihar |
| 24 | MS 32 | India | Tirupathi, Andhra Pradesh |
| 25 | MS 30 | India | Vallimalai, Tamil Nadu |
| 26 | MS 619 | India | Gorakhpur, Uttar Pradesh |
| 27 | JM 01 | India | - |
| 28 | JM 02 | India | - |
| 29 | MS 222 | India | Baripada, Odhisa |
| 30 | MS 608 | India | Aligarh, Uttar Pradesh |
| 31 | MS 611 | India | Kanpur, Uttar Pradesh |
| 32 | MS 108 | India | Mirzapur, Uttar Pradesh |
| 33 | MS 193 | India | Kanyakumari, Tamil Nadu |
| 34 | MS 287 | India | Amjheera, Madhya Pradesh |

| | | | |
|---|---|---|---|
| 35 | MS 509 | India | Amravati, Maharashta |
| 36 | MS 360.1 | India | Meleng, Assam |
| 37 | MS 401 | India | Sulur, Tamil Nadu |
| 38 | MS 629 | India | Barnpur, West Bengal |
| 39 | MS 327 | India | Kaundanyapur, Maharashtra |
| 40 | MS 518 | India | Bhivavaram, Andhra Pradesh |
| 41 | JM 05 | India | Bira, West Bengal |
| 42 | JM 06 | India | BR Hills, Karnataka |
| 43 | JM 08 | India | Agartala, Tripura |

**Table B2: Indoplanorbis sequences from previous studies**

| S.no. | Genbank accession number | Authors | Country | Region/state |
|---|---|---|---|---|
| 1 | GU451744 | Liu *et al* . (2010) | India | Assam |
| 2 | GU451745 | Liu *et al* . (2010) | Bangladesh | Mymensingh Division |
| 3 | GU451747 | Liu *et al* . (2010) | Indonesia | Java |
| 4 | GU451750 | Liu *et al* . (2010) | Laos | Vientiane |
| 5 | GU451746 | Liu *et al* . (2010) | Malaysia | Borneo |
| 6 | GU451738 | Liu *et al* . (2010) | Malaysia | Kampang Pelegong |
| 7 | GU451739 | Liu *et al* . (2010) | Nepal | Janakpur |
| 8 | GU451740 | Liu *et al* . (2010) | Oman | Wadi Bani Khaled |
| 9 | GU451741 | Liu *et al* . (2010) | Oman | Wadi Qab |
| 10 | GU451748 | Liu *et al* . (2010) | Philippines | Luzon |
| 11 | GU451742 | Liu *et al* . (2010) | Sri Lanka | Kekirawa |
| 12 | GU451743 | Liu *et al* . (2010) | Thailand | Khon Kaen |
| 13 | HM104223 | Liu *et al* . (2010) | Thailand | Phitsanulok |
| 14 | AY282587 | Albrecht *et al* . (2003) | Thailand | Loei Province |
| 15 | KR811332 | Devkota *et al* . (2015) | Nepal | Kamaraura, Dhanusa |
| 16 | KR811347 | Devkota *et al* . (2015) | Nepal | Tikauli, Chitwan |
| 17 | KR811348 | Devkota *et al* . (2015) | Nepal | Budhi Rapti, Chitwan |
| 18 | KR811338 | Devkota *et al* . (2015) | Nepal | Chitwan National Park |
| 19 | KR811349 | Devkota *et al* . (2015) | Nepal | Tikauli, Chitwan |
| 20 | KR811350 | Devkota *et al* . (2015) | Nepal | Dhumre river, Chitwan |
| 21 | KR811351 | Devkota *et al* . (2015) | Nepal | Tikauli, Chitwan |
| 22 | KR811352 | Devkota *et al* . (2015) | Nepal | KCF, Chitwan |

| | | | | |
|---|---|---|---|---|
| 23 | KR811339 | Devkota *et al* . (2015) | Nepal | Tikauli, Chitwan |
| 24 | KR811354 | Devkota *et al* . (2015) | Nepal | Tikauli, Chitwan |
| 25 | KR811353 | Devkota *et al* . (2015) | Nepal | KCF, Chitwan |
| 26 | KR811335 | Devkota *et al* . (2015) | Nepal | Khageri river, Chitwan |
| 27 | KR811336 | Devkota *et al* . (2015) | Nepal | Tikauli, Chitwan |
| 28 | KR811360 | Devkota *et al* . (2015) | Nepal | Chitwan National Park |
| 29 | KR811359 | Devkota *et al* . (2015) | Nepal | Chitwan National Park |
| 30 | KR811337 | Devkota *et al* . (2015) | Nepal | Chitwan National Park |
| 31 | KR811341 | Devkota *et al* . (2015) | Nepal | BCF, Chitwan |
| 32 | KR811342 | Devkota *et al* . (2015) | Nepal | BCF, Chitwan |
| 33 | KR811355 | Devkota *et al* . (2015) | Nepal | Sauraha, Chitwan |
| 34 | KR811333 | Devkota *et al* . (2015) | Nepal | Kamaraura, Dhanusa |
| 35 | KR811334 | Devkota *et al* . (2015) | Nepal | Kamaraura, Dhanusa |
| 36 | KR811356 | Devkota *et al* . (2015) | Nepal | Budhi Rapti, Chitwan |
| 37 | KR811357 | Devkota *et al* . (2015) | Nepal | Budhi Rapti, Chitwan |
| 38 | KR811358 | Devkota *et al* . (2015) | Nepal | Budhi Rapti, Chitwan |
| 39 | KR811344 | Devkota *et al* . (2015) | Nepal | Tulsi village, Dhanusa |
| 40 | KR811345 | Devkota *et al* . (2015) | Nepal | Tulsi village, Dhanusa |
| 41 | KR811346 | Devkota *et al* . (2015) | Nepal | Chisapani, Dhanusa |
| 42 | KR811361 | Devkota *et al* . (2015) | Nepal | Dhumre river, Chitwan |
| 43 | KR811340 | Devkota *et al* . (2015) | Nepal | Shishwar, Chitwan |
| 44 | KR811343 | Devkota *et al* . (2015) | Nepal | Chitwan National Park |
| 45 | KY024420 | Gauffre-Autelin *et al* .(2017) | Indonesia | South Sulawesi |
| 46 | KY024421 | Gauffre-Autelin *et al* .(2017) | Indonesia | South Sulawesi |
| 47 | KY024422 | Gauffre-Autelin *et al* .(2017) | Indonesia | Central Sulawesi |
| 48 | KY024423 | Gauffre-Autelin *et al* .(2017) | Indonesia | Central Sulawesi |
| 49 | KY024424 | Gauffre-Autelin *et al* .(2017) | Indonesia | Central Sulawesi |
| 50 | KY024425 | Gauffre-Autelin *et al* .(2017) | Indonesia | South Sulawesi |
| 51 | KY024426 | Gauffre-Autelin *et al* .(2017) | Indonesia | West Papua |
| 52 | KY024427 | Gauffre-Autelin *et al* .(2017) | Indonesia | West Java |
| 53 | KY024428 | Gauffre-Autelin *et al* .(2017) | Indonesia | West Java |
| 54 | KY024429 | Gauffre-Autelin *et al* .(2017) | Indonesia | West Papua |
| 55 | KY024430 | Gauffre-Autelin *et al* .(2017) | Indonesia | West Papua |
| 56 | KY024431 | Gauffre-Autelin *et al* .(2017) | Indonesia | West Papua |

| | | | | |
|---|---|---|---|---|
| 57 | KY024432 | Gauffre-Autelin *et al* .(2017) | Indonesia | West Papua |
| 58 | KY024433 | Gauffre-Autelin *et al* .(2017) | Indonesia | West Papua |
| 59 | KY024434 | Gauffre-Autelin *et al* .(2017) | Indonesia | West Papua |
| 60 | KY024435 | Gauffre-Autelin *et al* .(2017) | Indonesia | West Papua |
| 61 | KY024436 | Gauffre-Autelin *et al* .(2017) | Indonesia | West Papua |
| 62 | KY024437 | Gauffre-Autelin *et al* .(2017) | Indonesia | West Papua |
| 63 | KY024438 | Gauffre-Autelin *et al* .(2017) | Indonesia | West Papua |
| 64 | KY024439 | Gauffre-Autelin *et al* .(2017) | Indonesia | North Kalimantan |
| 65 | KY024440 | Gauffre-Autelin *et al* .(2017) | Indonesia | North Kalimantan |
| 66 | KY024441 | Gauffre-Autelin *et al* .(2017) | Indonesia | North Kalimantan |
| 67 | KY024442 | Gauffre-Autelin *et al* .(2017) | Indonesia | North Kalimantan |
| 68 | KY024443 | Gauffre-Autelin *et al* .(2017) | Indonesia | North Kalimantan |
| 69 | KY024444 | Gauffre-Autelin *et al* .(2017) | Indonesia | North Moluccas |
| 70 | KY024445 | Gauffre-Autelin *et al* .(2017) | Indonesia | North Moluccas |
| 71 | KY024446 | Gauffre-Autelin *et al* .(2017) | Indonesia | North Moluccas |
| 72 | KY024447 | Gauffre-Autelin *et al* .(2017) | Thailand | Phitsanulok |
| 73 | KY024448 | Gauffre-Autelin *et al* .(2017) | Thailand | Phitsanulok |
| 74 | KY024449 | Gauffre-Autelin *et al* .(2017) | Laos | Champasak |
| 75 | KY024450 | Gauffre-Autelin *et al* .(2017) | Laos | Champasak |
| 76 | KY024451 | Gauffre-Autelin *et al* .(2017) | Laos | Champasak |
| 77 | KY024452 | Gauffre-Autelin *et al* .(2017) | Myanmar | Kachin |
| 78 | KY024453 | Gauffre-Autelin *et al* .(2017) | Myanmar | Kachin |
| 79 | KY024454 | Gauffre-Autelin *et al* .(2017) | Myanmar | Kachin |
| 80 | KY024455 | Gauffre-Autelin *et al* .(2017) | Myanmar | Kachin |
| 81 | KY024456 | Gauffre-Autelin *et al* .(2017) | Indonesia | West Sulawesi |
| 82 | KY024457 | Gauffre-Autelin *et al* .(2017) | Indonesia | West Sulawesi |
| 83 | KY024458 | Gauffre-Autelin *et al* .(2017) | Indonesia | West Sulawesi |
| 84 | KY024459 | Gauffre-Autelin *et al* .(2017) | Indonesia | West Sulawesi |
| 85 | KY024460 | Gauffre-Autelin *et al* .(2017) | India | Assam |
| 86 | KY024461 | Gauffre-Autelin *et al* .(2017) | India | Tamil Nadu |
| 87 | KY024462 | Gauffre-Autelin *et al* .(2017) | India | Karnataka |
| 88 | KY024463 | Gauffre-Autelin *et al* .(2017) | India | Karnataka |
| 89 | KY024464 | Gauffre-Autelin *et al* .(2017) | India | Karnataka |
| 90 | KY024465 | Gauffre-Autelin *et al* .(2017) | India | Karnataka |

| | | | | |
|---|---|---|---|---|
| 91 | KY024466 | Gauffre-Autelin *et al* .(2017) | India | Karnataka |
| 92 | KY024467 | Gauffre-Autelin *et al* .(2017) | India | Karnataka |
| 93 | KY024468 | Gauffre-Autelin *et al* .(2017) | India | Kerala |
| 94 | KY024469 | Gauffre-Autelin *et al* .(2017) | India | Kerala |
| 95 | KY024470 | Gauffre-Autelin *et al* .(2017) | India | Tamil Nadu |
| 96 | KY024471 | Gauffre-Autelin *et al* .(2017) | India | Tamil Nadu |
| 97 | KY024472 | Gauffre-Autelin *et al* .(2017) | India | Karnataka |

**Table B3: Outgroup sequences**

| S.no. | Species | Species group | Genbank accession number (COI) | Genbank accession number (18S) | Genbank accession number (H3) |
|---|---|---|---|---|---|
| 1 | *Bulinus tropicus* | *B. tropicus* group | AM921842 | HM756332 | HM756451 |
| 2 | *Bulinus nyassanus* | *B. tropicus* group | AM921838 | HM756329 | HM756448 |
| 3 | *Bulinus natalensis* | *B. tropicus* group | AM286311 | HM756328 | HM756447 |
| 4 | *Bulinus forskalii* | *B. forskalii* group | AM286307 | HM756320 | HM756441 |
| 5 | *Bulinus wrighti* | *B. reticulatus* group | AM286318 | HM756323 | HM756444 |
| 6 | *Bulinus globosus* | *B. globosus* group | AM286290 | HM756309 | HM756428 |
| 7 | *Bulinus nasutus* | *B. globosus* group | AM286299 | HM756312 | HM756433 |
| 8 | *Bulinus obtusispira* | *B. globosus* group | | HM756313 | HM756434 |
